# Developmental Differences in Novelty Reactivity in Adolescent and Adult Male and Female Rats

**DOI:** 10.1101/2025.07.10.664152

**Authors:** Kirstie H. Stansfield, Antoniette M. Maldonado-Devincci, Cheryl L. Kirstein

**Author notes:** To whom correspondence should be addressed: Antoinette M. Maldonado-Devincci, PhD, Psychology Department-NSB 360, North Carolina A&T State University 1601 E. Market St, Greensboro, NC 27411.

## Abstract

Adolescence is a time of high-risk behavior and increased exploration. This developmental period is marked by a greater probability of initiating drug use and is associated with an increased risk to develop addiction and dependency in adulthood. Human adolescents are predisposed toward an increased likelihood of risk taking behaviors, including drug use or initiation. The purpose of the study was to examine differences in developmental risk-taking behaviors. Adolescent and adult animals were exposed to a novel stimulus in a familiar environment to assess impulsive behaviors, novelty preference and exploratory behaviors. Adolescent animals had greater novelty-induced locomotor activity, greater novelty preference, and showed higher approach and exploratory behaviors compared to preadolescent and adult animals. These data support the notion that adolescents may be predisposed toward sensation seeking and consequently are more likely to engage in risk taking behaviors, such as drug use initiation.

## I Introduction

Adolescence is a period when the brain is undergoing many complex changes that can exert long-term influences on decision making and cognitive processes (for review, see (Spear, 2000). It is also a period of experimentation, and Estroff and colleagues (1989) have reported that illicit drug use can begin as early as age 12, with peak periods of initiation between ages 15 and 19. The mean age of illicit drug initiation in adults categorized as having a substance use disorder is 16. /. years old, with initiation rare after age 20 (Anthony, 1991). In fact, initiation rates are so high that more than half (54%) of high school seniors have had at least one experience with an illicit compound [REFERENCE 70]. During the 1990’s, there was a steady rise in the frequency of cocaine use in teenagers, by 2003, 4.3% of eighth graders, 5.7% of tenth graders, and 8.2% of high school seniors reported frequent use of cocaine (Johnston, 2002). The fact that initiation of cocaine use is so dramatic during the adolescent period is particularly disconcerting given that the escalation of cocaine use appears more rapidly among teenagers than adult users, suggesting a greater addictive potential during adolescence than in adulthood (Estroff et al., 1989). Generally, adults who initiate drug use during adolescence are more likely to have higher lifetime rates of drug use and progress to dependency more rapidly than those who began drug use in adulthood (Clark, 1998; Helzer, 1991; Kandel et al., 1992). Moreover, adolescents demonstrate a more abrupt progression of illicit drug use and development of substance use disorders than adults (Warner et al., 1995), suggesting that this ontogenetic period renders the adolescent more vulnerable to addiction.

In humans, individual differences, including novelty-seeking are associated with alcohol and drug dependence. Novelty seeking and sweet liking are a combination of individual differences that predict alcoholic status both to a greater degree in men than women (Kampov-Polevoy et al., 2004). Novelty seeking alone was predictive of drug dependence status in both men and women (Kampov-Polevoy et al., 2004). During adolescence, sensation-seeking appears to be stable over at least 20 months in humans, and this personality trait appears to be highly correlated with drug use in teens (Pedersen, 1991). This is tempered by the specific construct assessed with disinhobition being the strongest predictor of drug-seeking behavior in human adolescents (Pedersen, 1991). Sex differences with experience seeking being correlated with cannabis use in males but not females during adolescence (Pedersen, 1991).

Adolescent males and females differ in their response to an inescapable novel environment, with adolescent males showing greater reactivity compared to adolescent females (Wooters et al., 2006). Adult females show greater novel environment-induced reactivity relative to their adult male counterparts (Wooters et al., 2006).

High responders as indexed by response to an inescapable novel environment show greater locomotor reactivity to d-amphetamine, however this relationship is tempered by dose, and is only observed at low doses (Kabbaj, 2006). Differences in novelty phenotype can differentially affect development of behavioral sensitization to d-amphetamine with high responders developing behavioral sensitization after a single administration of d-amphetamine and low responders developing behavioral sensitization after a number of injections of d-amphetamine (Kabbaj, 2006; Piazza et al, 1989). Novelty phenotype appears to affect acquisition of drug-taking behavior to a greater extent than maintenance of drug-taking behavior (Kabba, 2006). Others have shown high responding rats to develop sensitization to ethanol, but low respinding rats did not (Hoshaw and Lewis, 2001).

The frequency of substance use disorders is elevated in adults diagnosed with several psychological disorders (Anthony, 1991; Blanco et al., 2001; Bucholz, 1999; Helzer, 1991; Regier et al., 1990). Adolescents with similar disorders are also more likely to be diagnosed with substance use disorders (Shaffer et al., 2000; Swadi, 1999; Zeitlin, 1999). The fact that these mental disorders and adolescence are associated with substance use disorders suggests that common brain mechanisms may trigger drug susceptibility and potentially, addiction. These biological/neurochemical substrates might manifest into a behavioral trait or traits present in adolescents. Defective impulse control is a behavioral trait that characterizes psychiatric and substance use disorder groups (Moeller et al., 2001; Rogers and Robbins, 2001; Swadi, 1999). Adolescence is marked by high levels of risk taking behavior relative to individuals of other ages. Human adolescents exhibit a disproportional amount of reckless behavior, sensation seeking and risk taking (Trimpop, 1999). Not only is novelty seeking and high-risk behaviors during adolescence present in humans, but also non-human animals (Adriani et al., 1998; Douglas et al., 2003; Spear, 2000; Stansfield and Kirstein, 2006; Stansfield et al., 2004). Importantly, studies have demonstrated a strong correlation between novelty preference and impulsive reactivity with both the rewarding efficacy of psychomotor stimulants and self-administration rates in animals (Hooks et al., 1992; Klebaur et al., 2001). Researchers utilize two novelty preference paradigms: forced novelty exposure and free choice novelty exploration. Forced novelty exposure measures stress induced locomotor activity in a novel open field whereas free choice novelty exploration measures either frequency to approach a novel object or total time spent with a novel object in a familiarized environment (Stansfield and Kirstein, 2006). High responder (HR) adult rats to forced novelty show enhanced sensitivity to drug stimulant effects, higher rates of amphetamine and cocaine-induced locomotor activity and will self-administer these drugs more readily than low responder (LR) rats (Cools et al., 1997; Hooks et al., 1991; Piazza et al., 1989). Moreover, HR rats seem to participate in far greater risk taking behaviors and show much higher behavioral and neurochemical responses in reaction to environmental stressors or pharmacological challenges than LR rats (Bevins, 1997; Klebaur et al., 2001). Pelloux et al. demonstrated that HR to forced novelty exposure consumed less oral amphetamine compared to LR (Pelloux et al., 2004). Additionally, Pelloux and colleagues demonstrated that novelty preference is positively correlated with consumption of a low concentration morphine solution (Pelloux et al., 2006). A modest relationship was observed in adolescent male and female mice for high novelty mice to consume more nicotine as compared to low novelty mice (Abreu-Villaca et al., 2006). Novel environment-induced reactivity was a strong predictor of subsequent responsivity to repeated methylphenidate exposure (Wooters et al., 2006). This effect was greatest in adult females compared to adult males and adolescent-females (Wooters et al., 2006). Taken together, these data suggest a relationship between novelty-seeking and drug use, making it more likely that adolescent’s will become involved in risky behaviors which may include drug use, initiation and increased vulnerability to the rewarding properties of these drugs.

No sex differences in novelty preference, defined by number of head dips in a hole board task, or anxiety, defined as the percentage of center squared crossed in the hole board task, were observed in early adolescent male and female mice (Abreu-Villaca et al., 2006). In humans, men were almost two times more likely to be classified as an alcoholic compared to women ((Kampov-Polevoy et al., 2004). Sex differences in ambulation have been observed in adulthood, however this effect was not observed at PND 35 or PND 45 (Masur et al., 1980).

In humans, novelty seeking scores were predictive of drug dependence, but not alcoholic status (Kampov-Polevoy et al., 2004).

The aim of this study was investigate novelty preference behaviors across development between female and male animals. For this purpose, the current study compares reactivity to novelty using both forced novelty exposure and free choice novelty preference and exploration.

## II. Methods

One hundred male and female Sprague-Dawley (Harlan Laboratories, Indianapolis, IN) rats, offspring of established breeding pairs in the laboratory (University of South Florida, Tampa) were postnatal day (PND) 24 [periadolescent], PND 34 [early adolescent], PND 44 [late adolescent] and PND 59 [young adult] at the time of testing. No more than one rat per litter per sex or per age was used in a given condition. Pups were sexed and culled to 10 pups per litter on PND 1. Pups remained housed with their respective dams in a temperature and humidity-controlled vivarium on a 12:12 h light:dark cycle (07:00 h/19:00 h) until PND 21, following which pups were weaned and group housed for the remainder of the exeriment. Animals were naïve/ unmanipulated until the time of training. The care and use of animals was in accordance with local standards set by the Institutional Animal Care and Use Committee and the NIH Guide for the Care and Use of Laboratory Animals (National Institutes of Health, 1986).

### Procedure

Animals were tested on a black plastic circular platform (116 cm in diameter) standing 70 cm from the ground, with a white plastic barrier (48 cm in height) enclosing the arena (100 cm in diameter). A video camera was suspended directly over the table and recorded the animal’s behavior using a Noldus Behavioral Tracking System.

Over a period of four consecutive days, each rat (PND 21-24, 31-34, 41-44 and 56-59) was placed on the open field in one of four randomly selected zones and allowed to freely explore the novel environment for five minutes. This procedure was performed twice a day for a total of 8 habituation trials. Immediately following the 8^th^ trial, animals were removed for 1 minute while a single novel object (approximately 6.5 in. high) was attached to the center of the table (trial 9). Rats were placed in a random zone and allowed to explore the familiar environment for five minutes. Forced-novelty locomotor activity (total distance moved on trial 1), free choice novelty preference (time spent with novel object in familiarized environment), and free choice novelty exploration (frequency to approach novel object in familiarized enviroment).

### Data Analyses

Data analyses were performed with Graphpad Prism (Graphpad). The data were expressed as the means +-SEM, and the significance level was set at p=0.05. Separate one-way analyses of variance (ANOVA) were performed on forced-novelty locomotor activity, free choice novelty preference and free choice novelty exploration with subsequent post hoc analyses (Newman-Keuls Multiple Comparison Test) to isolate differences across age or sex.

## Results

### Forced novelty locomotor activity

The present findings demonstrate that both male and female preadolescent rats exhibited decreased forced choice locomotor activity compared to all other ages and sex [F(3,104)=70.70, p<0.0001, see Figure 1]. When initially placed in the apparatus, preadolescent animals were more reactive, whether it be stress-induced or novelty-induced exploration, than early-adolescent, late adolescent and young adult rats.

**Figure 1.**
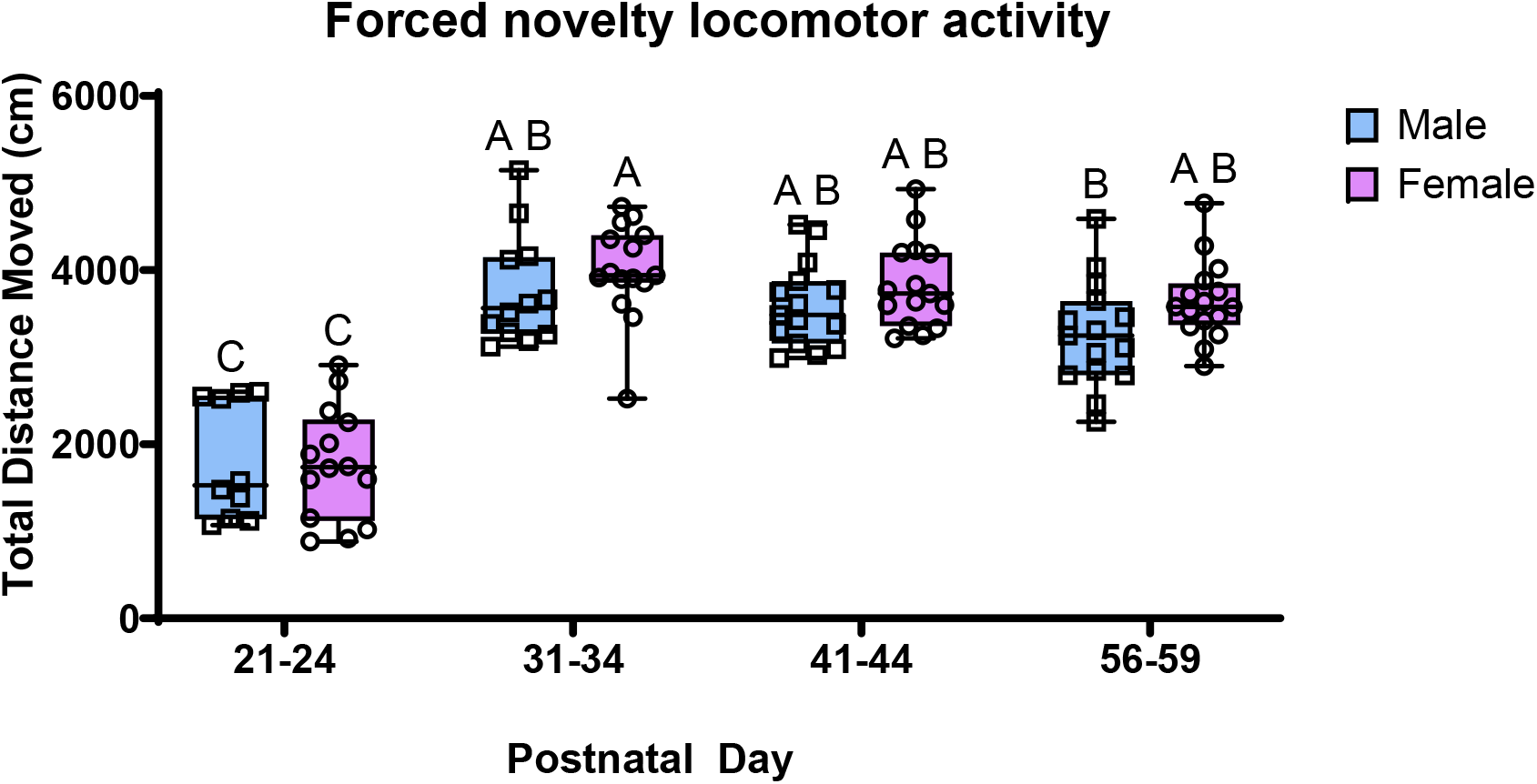

### Free-choice novelty preference

A one-way ANOVA revealed significant differences between free-choice novelty preference, age and sex [F(3, 99)=6.4,p,0.005]. Preadolescent female rats differed significantly from both late-adolescent male and female and young adult male rats. In addition, early adolescent male rats differed significantly from late-adolescent female and young adult male rats (see Figure 2). Preadolescent females demonstrate greater preference for the novel object compared to any other female age group. Moreover, early adolescent males demonstrate greater preference for the novel object compared to late adolescent females and young adult male rats. Overall, there seems to be a trend for increased novelty preference during periadolescent that decreases across age into young adulthood.

**Figure 2.**
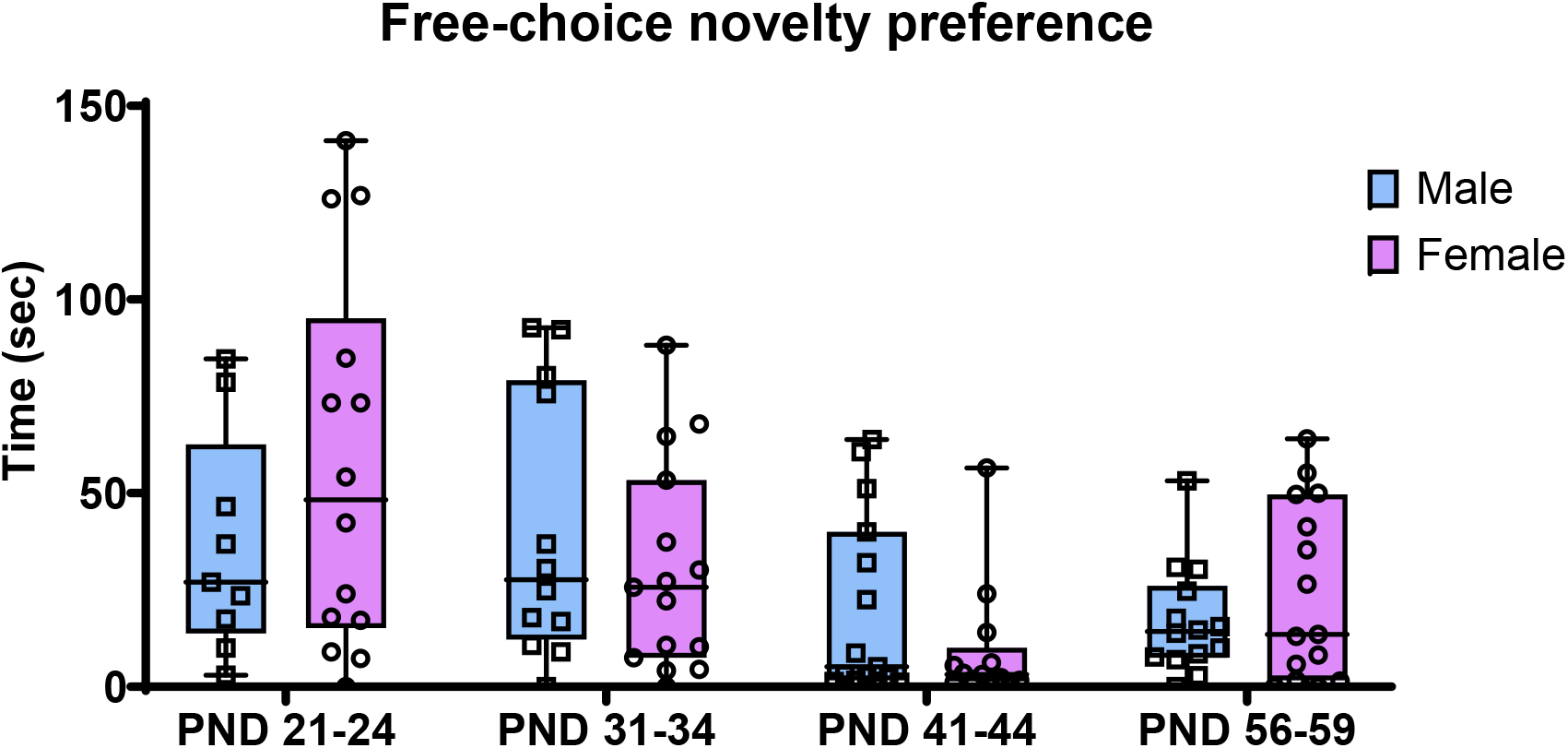

### Free-choice novelty exploration

The present findings demonstrate that early adolescent male rats differed from late adolescent female and young adult male rats [F(3, 95)=5.4, p<0.01, see Figure 3]. Early adolescent males approached the novel object significantly more than late adolescent female and young adult male rats.

**Figure 3.**
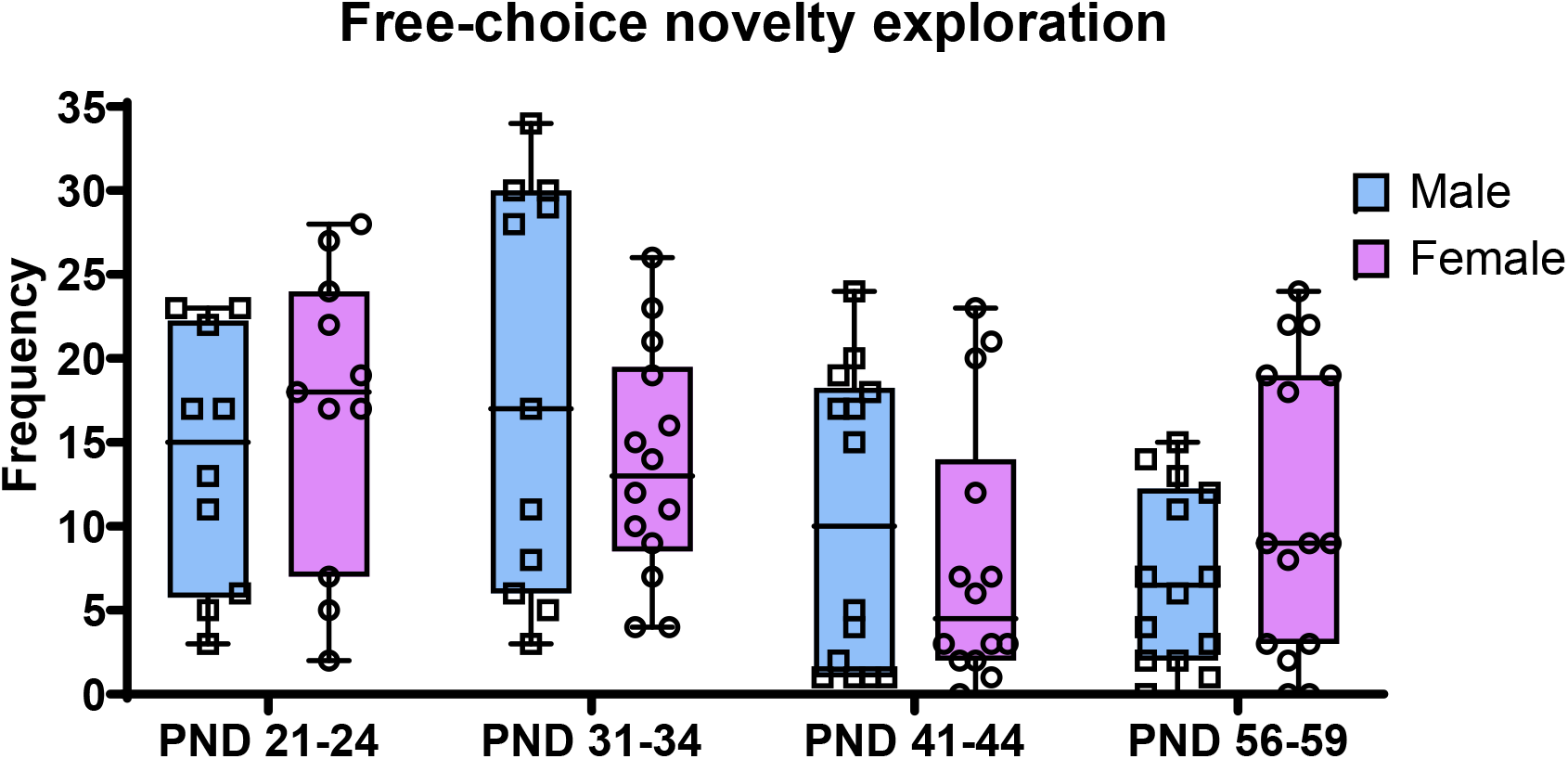

## Discussion

Previous work in adult animals has demonstrated that a preference for novelty is indicative of a facilitated acquisition of drug use (Bevins, 1997). Novelty preference has typically been defined as increased locomotor activity in a novel environment. Adolescent animals and humans who prefer novelty are more likely to use/ abuse drugs and individuals who initiate use in adolescence will progress to dependency more rapidly than those who began drug use in adulthood (Clark, 1998; Helzer, 1991; Kandel et al., 1992). The aim of the present study was to investigate developmental and sex differences in response to both forced novelty exposure and free choice novelty.

The present data provide evidence that periadolescent animals were more reactive to forced novelty than early-adolescent, late adolescent and young adult rats. This behavioral characteristic appears to be transient across development, with an increase during periadolescent and a subsequent decrease across adolescence into young adulthood. In addition, periadolescent females demonstrate greater preference for the novel object compared to any other female age group and compared to young adult male rats. Moreover, early adolescent males demonstrate greater preference for the novel object compared to late adolescent females and young adult male rats. Overall, there seems to be a trend for increased novelty preference during periadolescent that decreases across time into young adulthood. Finally, early adolescent males approached the novel object significantly more than late adolescent female and young adult male rats. Novelty-seeking behaviors are an innate behavior in both human and non-human adolescents. Importantly, studies have demonstrated a strong correlation between novelty preference and the rewarding efficacy of psychomotor stimulants and self-administration rates in animals (Hooks et al., 1992; Klebaur et al., 2001). High novelty seeking rats show higher rates of amphetamine and cocaine-induced locomotor activity and will self-administer these drugs more readily than low novelty seeking rats (Hooks et al., 1992). Moreover, high novelty seeking rats seem to participate in far greater risk-taking behaviors and show much higher behavioral and neurochemical responses in reaction to environmental stressors or pharmacological challenges than low novelty seeking rats (Bevins, 1997; Klebaur et al., 2001). Additionally, adolescent animals classified as high responders to novelty based on activity in a novel environment and also by time spent with a novel object in a familiar environment exhibited greater morphine place conditioning in adulthood compared to low responders to novelty (Zheng et al., 2004). The present data provide evidence depending on the novelty preference measure (i.e forced-exposure novelty vs. free-choice novelty preference/ exploration) that young adolescents are at greater risk to engage in drug use that may predisopose them to continued use rats.

Similar to previous work, adolescent rats spend significantly more time with the novel object as compared to adult rats (Douglas et al., 2003). Additionally, no within-age sex differences were observed in the present experiment, similar to previous work (Douglas et al., 2003). When animals were separated into novelty phenotype based on reactivity to a novel environment, adult rats exhibited greater novelty scores relative to adolescents, and adult females had greater scores than adult males and adolescent males had greater scores than adolescent females (Wooters et al., 2006). In contrast, when animals were separated based on a novelty place preference test, there were no age or sex differences in novelty response in any of the groups (Wooters et al, 2006). These data are suggestive of a difference in these two constructs and each may be differentially indicative of response to drugs of abuse.

No sex differences in nicotine consumption were present in high novelty and low novelty or high anxiety and low anxiety adolescent mice (Abreu-Villaca et al., 2006). On three (PND 35-37) of ten (PND 31-40) days of adolescent exposure to a nicotine solution in a two-bottle choice paradigm, high novelty mice consumed significantly more nicotine relative to low novelty mice (Abreu-Villaca et al., 2006). No significant differences in nicotine consumption were observed between high anxiety and low anxiety adolescent male and female mice (Abreu-Villaca et al., 2006).

Sex differences in ambulation scores were observed at PND 60 in male and female adult rats (Masur et al., 1980). A trend for a similar effect was observed in the present experiment for Total Distance Moved on Trial 1. This effect likely did not reach statistical significance in the present experiment given different animals were used at each age in the present experiment and the same animals were used across age in the experiment by Masur and colleagues.

These data suggest that both periadolescent male and female rats are more at risk to engage in drug use. Importantly, greater activity in a novel environment may be indicative of increased stress or anxiety or enhanced neophobia. Van den Buuse et. al (2001) have demonstrated that exposure to the novelty of an open field causes an increase in blood pressure, heart rate, body temperature and exploratory locomotor activity, results indicate that an increase in locomotor activity in a novel environment is stressful or anxiogenic. Together, these data show that adolescent rats are significantly more reactive to and prefer novel stimuli. There appears to be an initial sex-dependent difference in novelty preference in adolescent rats, with preadolescent females exhibiting increased novelty preference. Therefore, these data would suggest the notion that preadolescent females may be more sensitive to this behavioral characteristic affecting subsequent behaviors, such as responsivity to drugs. Young adolescent males that show high novelty preference subsequently may be more susceptible to the effects of drugs of abuse.

Sex differences during adolescence moderated differences in responsivity to repeated methylphenidate during adolescence (Wooters et al., 2006). Specifically, high responding adolescent males were more sensitive to 3 mg/kg repeated methylphenidate compared to their adolescent female counterparts (Wooters et al., 2006). These high responding adolescent males show a similar behavioral profile as compared to high responding adult females, indicating that both age and sex can alter the effects of novelty phenotype on responsivity to repeated methylphenidate (Wooters et al., 2006).

Interestingly, when animals were separated based on free-choice novelty as indexed by time spent in a novel compartment relative to a familiar compartment, this did not predict locomotor reactivity to repeated methylphenidate (Wooters et al., 2006).

It seems that a dissociation exists between forced novelty exposure and free choice novelty exploration in adolescent rats, suggesting that stress-induced locomotion and novelty-seeking behavior are different biobehavioral phenomena and might be activated by different neural and hormonal substrates. Interestingly, the relationship between free choice novelty exploration and cocaine place conditioning differs between adolescent and adult rats suggesting individual differences in free choice novelty exploration may be an important behavioral characteristic that predisposes adolescents to engage in cocaine use and demonstrate increased vulnerability to drug dependence.

